# StrIDR: a database of intrinsically disordered regions of proteins with experimentally resolved structures

**DOI:** 10.1101/2024.08.22.609111

**Authors:** Kartik Majila, Shruthi Viswanath

## Abstract

**Motivation:** Intrinsically disordered regions (IDRs) of proteins exist as an ensemble of conformations, and not as a single structure. Existing databases contain extensive, experimentally derived annotations of intrinsic disorder for millions of proteins at the sequence level. However, only a tiny fraction of these IDRs are associated with an experimentally determined protein structure. Moreover, even if a structure exists, parts of the disordered regions may still be unresolved.

**Results:** Here we organize **Str**uctures of **I**ntrinsically **D**isordered **R**egions (StrIDR), a database of IDRs confirmed *via* experimental or homology-based evidence, resolved in experimentally determined structures. The database can provide useful insights into the dynamics, folding, and interactions of IDRs. It can also facilitate computational studies on IDRs, such as those using molecular dynamics simulations and/or machine learning.

**Availability:** StrIDR is available at https://isblab.ncbs.res.in/stridr. The web UI allows for downloading PDB structures and SIFTS mappings of individual entries. Additionally, the entire database can be downloaded in a JSON format. The source code for creating and updating the database is available at https://github.com/isblab/stridr.

## Introduction

Intrinsically Disordered Proteins (IDPs) or Intrinsically Disordered Regions (IDRs) of proteins are characterized by a lack of well-defined three-dimensional structures in their monomeric form; instead, they exist as an ensemble of conformations (Dunker et al., 2002; Oldfield & Dunker, 2014; Uversky, 2013). Herein, we use IDR to refer to both of the above. IDRs constitute a significant portion, approximately 33-50% of the eukaryotic proteome (McConnell & Parker, 2023; Oldfield & Dunker, 2014; Vucetic et al., 2003). IDRs play diverse roles in the cell, including molecular recognition, signal transduction, intracellular transport, scaffolding, chaperoning protein folding, and condensate formation (Fonin et al., 2018; Uversky, 2013). Their intrinsic disorder enables them to possess a large capture radius, overcome steric restrictions, form transient and promiscuous interactions/interact transiently and with multiple partners, and undergo induced folding (Fonin et al., 2018). Upon binding to a partner, IDRs may attain an ordered structure (disorder-to-order or DOR transition) or remain disordered (disorder-to-disorder or DDR transition) (Miskei et al., 2020). An estimated 20% of IDRs in eukaryotic proteomes fold upon binding and are associated with a high prevalence of disease mutations (Alderson et al., 2022).

Given their importance for biological function, IDPs have been extensively studied in isolation and in complex with a partner protein. This data has been collected in various IDR-specific databases (Table S1) (Aspromonte et al., 2024; Di Domenico et al., 2012; Fichó et al., 2017; Fukuchi et al., 2012, 2014; Miskei et al., 2017, 2020; Piovesan et al., 2017, 2018, 2021, 2022; Schad et al., 2018; Sickmeier et al., 2007). Disprot and MobiDB are among the most comprehensive databases on IDRs (Aspromonte et al., 2024; Di Domenico et al., 2012; Piovesan et al., 2017, 2018, 2021, 2022; Vucetic et al., 2005). Disprot provides the most extensive manual curation of experimental data on the structures and functions of IDRs (Aspromonte et al., 2024; Piovesan et al., 2017; Vucetic et al., 2005). MobiDB combines data from several databases and prediction methods to provide more complete disorder annotations for protein sequences along with information about the structure, function, and dynamics of IDRs (Di Domenico et al., 2012; Piovesan et al., 2018, 2021). The data in MobiDB is a combination of manually curated annotations from experimental data, such as those from Disprot and IDEAL, extensions of these annotations to homologous proteins, inferences on disordered and binding residues based on structures using methods such as FLIPPER and Mobi, as well as sequence-based predictions. Importantly, it allows the user to choose the annotations based on the quality of data aggregated to form the annotation.

Several databases provide information about the interactions of IDRs. ELM provides annotations of Eukaryotic Linear Motifs (ELM) and Short Linear Motifs (SLiMs), whereas DisBind is dedicated to the classification of IDR binding sites to partner proteins, DNA, RNA, and metal ions (Kumar et al., 2022; Puntervoll et al., 2003; Yu et al., 2017). IDEAL is a manually curated database on the structure, binding, and post-translational modifications (PTM) of experimentally verified IDRs. It also provides assignments of structural domains, focusing on protean segments (ProS) – IDRs which undergo coupled folding and binding (Fukuchi et al., 2012, 2014). DIBS and MFIB consist of IDRs in complexes where the IDR undergoes a DOR transition (Fichó et al., 2017; Schad et al., 2018). The former includes complexes formed by IDRs upon binding to an ordered partner whereas the latter comprises complexes formed exclusively by IDRs. Finally, FuzDB curates complexes where the IDR undergoes a DDR transition (Miskei et al., 2017).

Although the databases mentioned above contain millions of sequences containing IDRs, a tiny fraction of the IDRs in these sequences are associated with an experimentally determined protein structure. Moreover, a disordered region of interest could be unresolved in the structure even if associated with one. The number of these IDRs that are resolved in experimental structures is unknown. Here, we organize **Str**uctures of **I**ntrinsically **D**isordered **R**egions (StrIDR), a database of protein regions verified to be IDRs *via* experiments or homology-based evidence, that are resolved in experimentally determined protein structures. It includes PDB entries with IDR containing at least one disordered residue resolved in the structure. We identify 84596 PDB entries containing an IDR sequence, of which 72001 contain IDRs resolved in the structure (Fig. 1).

**Figure 1:**
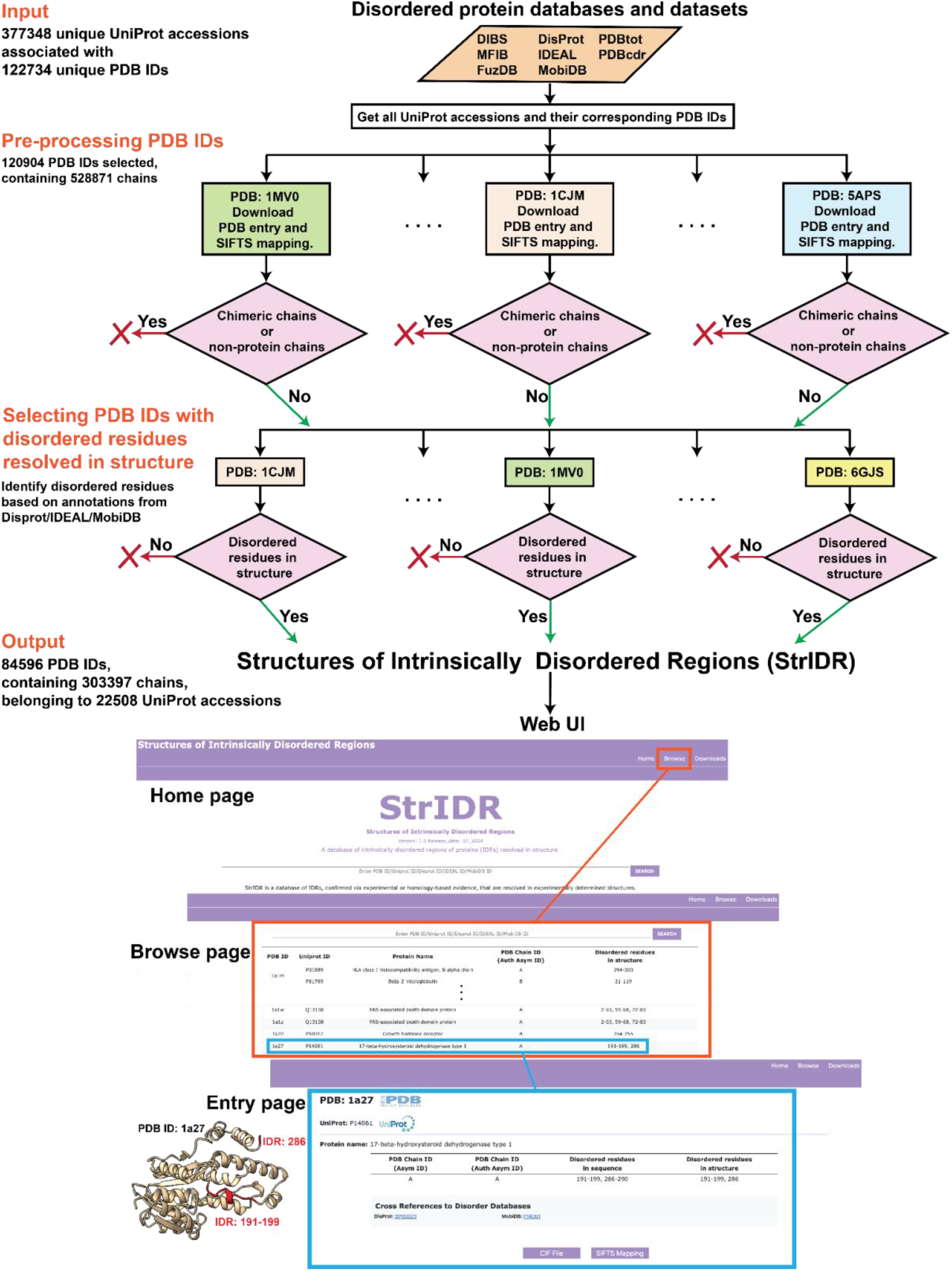
StrIDR creation workflow and web UI. The StrIDR database incorporates information gathered from disordered protein databases and datasets including DIBS, MFIB, FuzDB, DisProt, IDEAL, MobiDB, PDBTot, and PDBCDR. The inputs from these databases and datasets include the UniProt accessions of IDRs. The PDB entries corresponding to the IDR UniProt accessions are retrieved and processed. Subsequently, the PDB entries containing disordered residues in the structure are identified based on the disorder annotations from DisProt/IDEAL/MobiDB. The StrIDR web UI comprises Home, Browse, and Download pages. The StrIDR entry on PDB 1a27 chain A has been shown as an example.

StrIDR is expected to be useful for researchers who seek to gain insights into the dynamics, folding, and interactions of IDRs. It would facilitate a detailed analysis of the conformational flexibility and structural heterogeneity of IDRs, identifying patterns in flanking regions, and the elucidation of the interactions of IDRs with their molecular partners. This would lead to a deeper understanding of the biological roles of IDRs and their involvement in various cellular processes. Further, it may enable the identification of potential drug targets of IDRs. Finally, StrIDR can also be useful for methods developers by providing data for training and validating computational methods for studying IDRs, such as molecular dynamics (MD) simulations and machine learning (ML) methods (Lindorff-Larsen & Kragelund, 2021).

## Database Construction

### Gathering data from disordered protein databases

First, we gathered the available UniProt accessions for proteins containing IDRs from databases of disordered proteins: DIBS (Fukuchi et al., 2014), MFIB (Fichó et al., 2017), FuzDB (Miskei et al., 2017), DisProt (Aspromonte et al., 2024; Piovesan et al., 2017; Vucetic et al., 2005), IDEAL (Fukuchi et al., 2012, 2014), and MobiDB (Di Domenico et al., 2012; Piovesan et al., 2018, 2021) (Fig. 1). From IDEAL, we obtained UniProt IDs for entries annotated as “verified ProS”, *i*.*e*., sequences that have been experimentally verified to be disordered in isolation and ordered upon binding (Fukuchi et al., 2012, 2014). From MobiDB, we obtained UniProt accessions for entries annotated with “curated-disorder-priority”, “homology-disorder-priority”, “curated-lip-priority”, “homology-lip-priority”,”derived-lip-priority”, “derived-binding_mode_context_dependent-priority”, “curated-binding_mode_disorder_to_disorder-priority”, “derived-binding_mode_disorder_to_disorder-priority”, “homology-binding_mode_disorder_to_disorder-priority”, “derived-binding_mode_disorder_to_order-priority”, “derived-mobile-th_90”, “derived-mobile context dependent-th_90” (Di Domenico et al., 2012; Piovesan et al., 2018, 2021). These annotations refer to direct and derived experimental or homology-based evidence for disorder.

We also included Uniprot accessions from the PDBtot and PDBcdr datasets (Miskei et al., 2020). PDBtot consists of IDRs that either undergo a DOR or DDR transition, whereas PDBcdr consists of IDRs that undergo both DOR and DDR transitions depending upon the partner. In total, 377348 unique Uniprot accessions of IDRs were obtained.

Next, the PDB REST API was queried to obtain all the PDB entries associated with the given Uniprot accessions, along with entry and entity-level information for each PDB entry (Berman et al., 2000; Burley et al., 2022). This resulted in a total of 122734 unique PDB entries (Table S1).

### Pre-processing PDB entries

The PDB entries were downloaded using the Python requests library. PDB entries were mapped to UniProt sequences using the Structure Integration with Function, Taxonomy, and Sequence (SIFTS) tool (Dana et al., 2019; Velankar et al., 2013). Subsequently, entries with obsolete PDB IDs were removed and those with deprecated PDB IDs were replaced with the corresponding superseding entries. PDB entries without a SIFTS mapping were excluded. Finally, chimeric and non-protein entities were removed from each PDB entry (Fig. 1).

### Selecting PDB entries with disordered residues resolved in the structures

Next, for each PDB entry, we identified the disordered residues in a chain. This was achieved by cross-referencing the UniProt mapping of the chain with the annotations in DisProt, IDEAL, and MobiDB. Missing residues, *i*.*e*., residues that are not resolved in the structure, were then removed from the chain. Chains with at least one disordered residue resolved in the structure were selected (termed “IDR chains”). PDBs with at least one such chain were retained. This resulted in a total of 84596 PDB entries with 303397 IDR chains corresponding to 22508 UniProt accessions (Fig. 1). 72001 of these PDB entries contain 145659 IDR chains with five or more disordered residues resolved in the structure (Fig. S1).

### Database Implementation and Maintenance

StrIDR has been developed with the Django framework and the data is stored using the SQLite database engine. The front end was implemented using HTML, CSS, and Bootstrap 5.0. The implementation is modular and designed for ease of maintenance.

### Database User Interface and Example

All entries in StrIDR are available through a dedicated web server (Fig. 1). The “Home” page describes the purpose of the database to new users and provides statistics on the database. The “Browse page” lists the entries in the database in a tabular layout. Each entry in StrIDR is identified by its PDB ID, for which more information can be found on the entry page. The entry page includes all the chains in the PDB that have disordered residues resolved in the structure. For each such chain, we provide the associated UniProt accessions, the protein name, and a table containing detailed information about the chain. The “Asym ID” and “Auth Asym ID” refer to the PDB-assigned, and the author-assigned chain IDs respectively. “Disordered residues in sequence” displays all contiguous stretches of UniProt residue positions annotated as disordered in the chain. Similarly, “Disordered residues in structure” displays the UniProt residue positions for all contiguous stretches of disordered residues present in the structure. Cross-references to the database from which the disorder annotation was obtained are provided for each UniProt accession. Finally, we provide the PDB structure of an entry in the CIF file format and its SIFTS mapping in a TSV file format.

StrIDR allows querying for entries based on their PDB, UniProt, DisProt, IDEAL, and MobiDB accessions. This search functionality is accessible from both the Home and Browse pages. Additionally, the complete database can be downloaded from the Downloads page, including the UniProt sequences in JSON format, as well as the CIF files and SIFTS mappings for all StrIDR entries.

Fig. 1 shows an example of a StrIDR entry for PDB ID 1a27 for which the structure was determined *via* X-ray crystallography. It contains a single chain corresponding to 17-beta-hydroxysteroid dehydrogenase type 1, an alcohol oxidoreductase involved in testosterone synthesis (Mazza et al., 1998). The PDB entry includes residues 2-289. Disorder annotations from DisProt and MobiDB highlight the residues 191-199 and 286-290 to be IDRs due to the electron density being less well-defined in X-ray structures containing this region (Ghosh et al., 1995; Mazza et al., 1998). However, only IDR residues 191-199 and 286 are resolved in the structure.

In summary, StrIDR provides a comprehensive resource for IDRs resolved in experimentally determined structures. It integrates information from several IDR-related structure and sequence databases including DIBS, MFIB, FuzDB, DisProt, IDEAL, and MobiDB, along with datasets of disordered proteins in complexes, such as PDBtot and PDBcdr. Despite efforts such as StrIDR, there is still a need for a concerted community-wide effort to obtain more structural data on IDRs (Jahn et al., 2024; Piovesan et al., 2022). This would not only further our understanding of the roles of IDRs in processes such as regulation and signaling, but also enable the development of novel computational methods, such as deep generative models for IDRs.

## Data availability

The database is available at https://isblab.ncbs.res.in/stridr. Files containing the input data and scripts are available at GitHub https://github.com/isblab/stridr.

## Acknowledgment

We thank lab members Shreyas Arvindekar, Muskaan Jindal, Omkar Golatkar, and Mubashira KP for their useful comments on the manuscript. Molecular graphics images were produced using the UCSF Chimera and UCSF ChimeraX packages from the Resource for Biocomputing, Visualization, and Informatics at the University of California, San Francisco (supported by NIH P41 RR001081, NIH R01-GM129325, and National Institute of Allergy and Infectious Diseases).

