## Supplementary Information for "StrIDR: a database of intrinsically disordered regions of proteins with experimentally resolved structures"

### Supplementary Figures

**Figure S1: Distribution of disordered residues in the StrIDR database.** Number of disordered residues per PDB entry included in StrIDR. For ease of visualization, the maximum number of disordered residues per PDB has been capped to 1000.

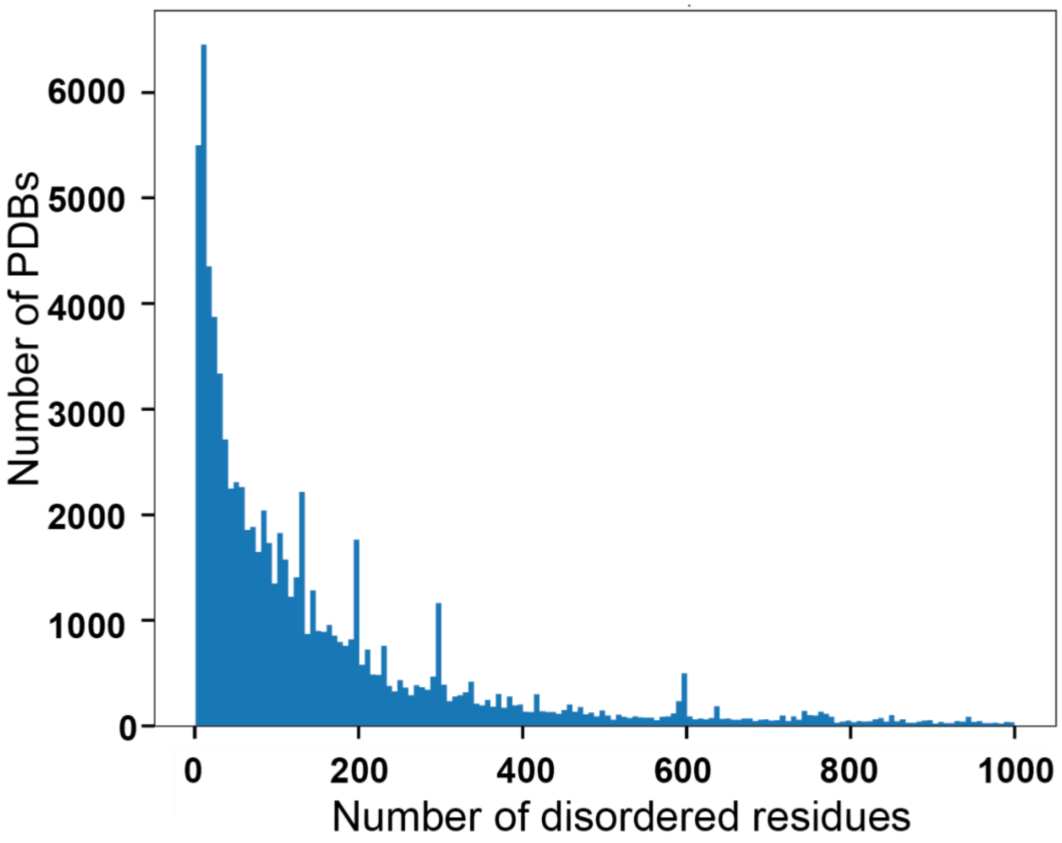

### Supplementary tables

**Table 1. Database Sources.** Disordered protein databases and datasets used for StrIDR database creation. We obtain UniProt entries from all the databases. The acronyms used are as follows: IDR: intrinsically disordered region of a protein, DOR: disorder-to-order, DDR: disorder-to-disorder.

| Database | Description | URL | Reference | Total UniProt IDs |
|---|---|---|---|---|
| DIBS | Database of complexes between an IDR and an ordered partner protein; the IDRs undergo a DOR transition | https://dibs.enzim.ttk.mta.hu/ | (Schad et al., 2018) | 772 |
| MFIB | Database of complexes between IDRs; the IDRs undergo a DOR transition | https://mfib.enzim.ttk.mta.hu/ | (Fichó et al., 2017) | 1329 |
| FuzDB | Dataset of complexes containing IDRs; the IDR undergoes a DDR transition | https://fuzdb.org/ | (Miskei et al., 2017) | 404 |
| DisProt | Manually curated repository for structure and function of experimentally verified IDRs. | https://disprot.org | (Aspromonte et al., 2024) (Piovesan et al., 2017) (Vucetic et al., 2005) | 3064 |
| IDEAL | Database for experimentally verified IDRs with focus on Protean segments. | https://www.ideal-db.org/ | (Fukuchi et al., 2014) (Fukuchi et al., 2012) | 1015 |
| MobiDB | Database for predictions and annotations on protein disorder and mobility | https://mobidb.bio.unipd.it/ | (Piovesan et al., 2023) (Piovesan et al., 2018) (Di Domenico et al., 2012) | 375909 |

| | | | | |
|---|---|---|---|---|
| | | | (Piovesan et al., 2017) 22-08-2024 09:15:00 | |
| PDBTot | Dataset of complexes containing IDRs; the IDR undergoes either a DOR or a DDR transition | - | (Horvath et al., 2020)<br><br>(Miskei et al., 2020) | 513 |
| PDBCDR | Dataset of complexes containing IDRs; the IDR undergoes both DOR and DDR transitions in a context-dependent manner | - | (Horvath et al., 2020)<br><br>(Miskei et al., 2020) | 164 |
